## Supplementary Information for "Transmission of SARS-CoV-2 in domestic cats imposes a narrow bottleneck"

### Supplementary Figures

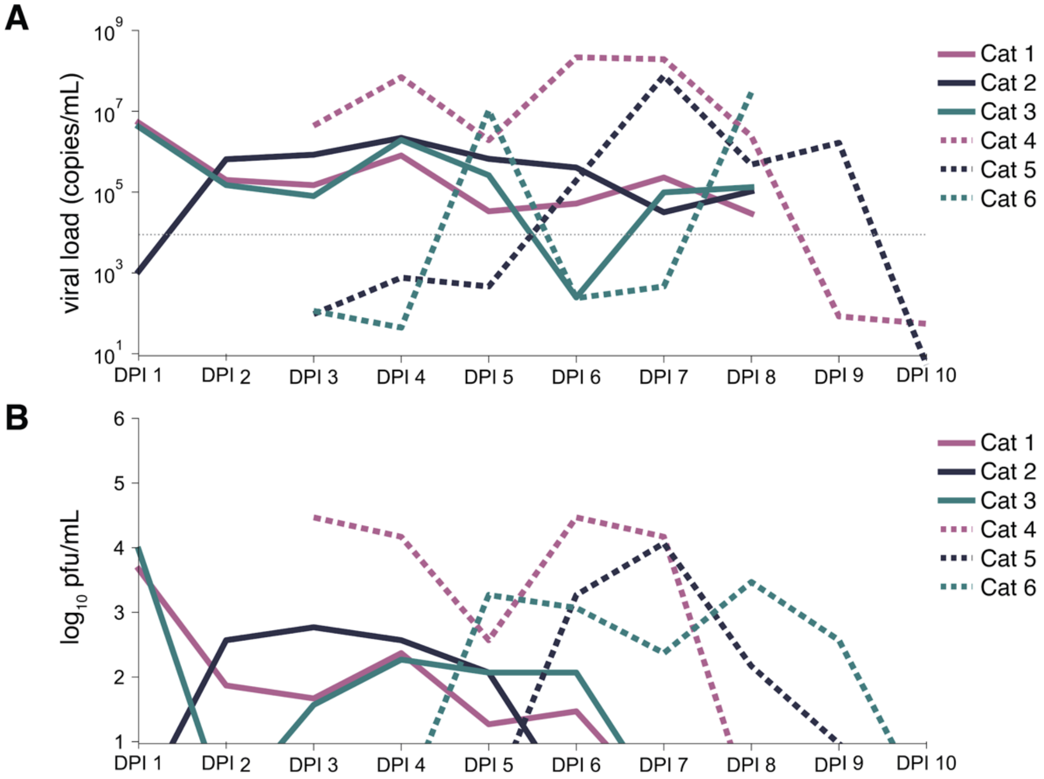


**Supplementary Figure 1**. **Viral loads and viral titers over time**. A) Viral RNA burden over time for each cat. Index cats are represented by a solid line and contact cats are represented by a dashed line. Transmission pairs are denoted by color. The grey, horizontal dotted line represents when less than ~100 copies/μL are input into the reverse transcription reaction. B) Infectious viral titer over time. Index cats are represented by a solid line and contact cats are represented by a dashed line. Transmission pairs are denoted by color.

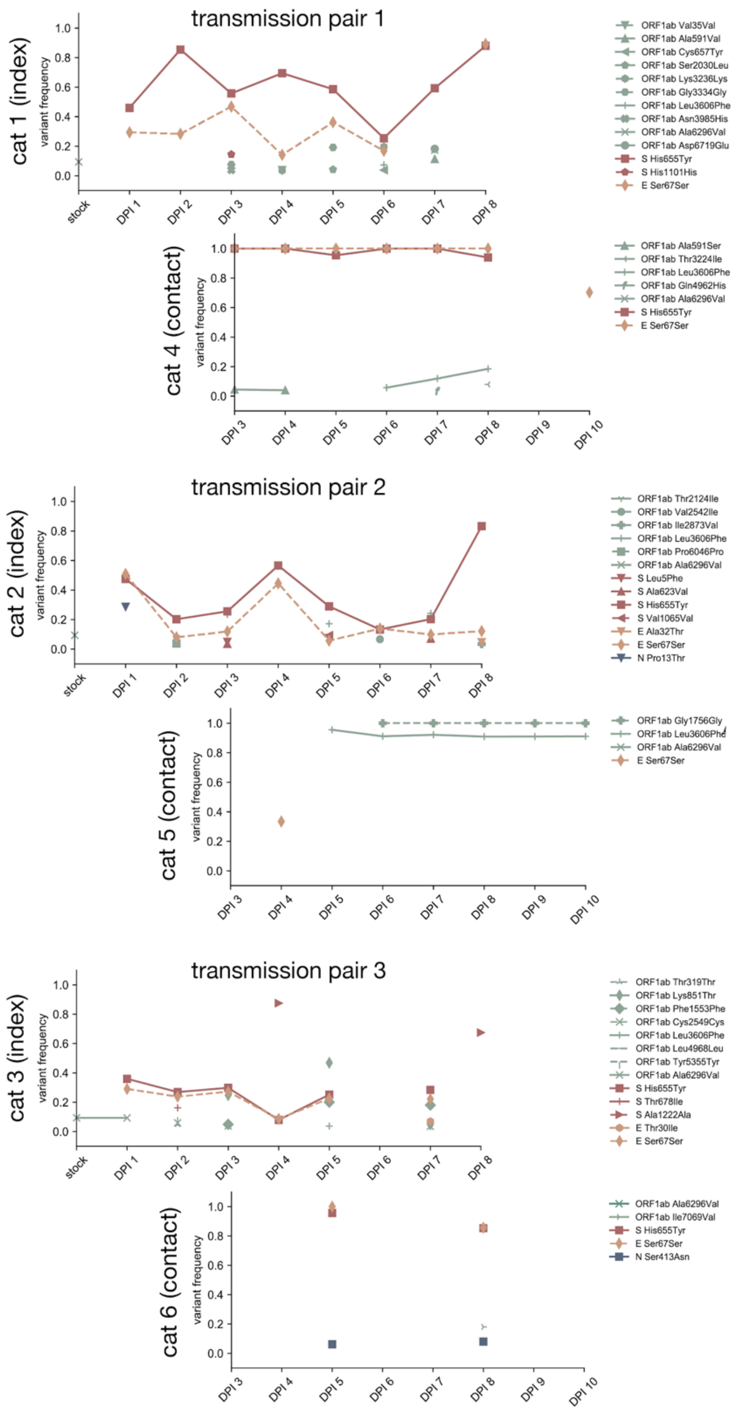


**Supplementary Figure 2. Longitudinal frequency of iSNVs detected in all cats and at all timepoints.** Each variant is colored based on gene location. Nonsynonymous variants are plotted with solid lines and synonymous variants are plotted with dashed lines. Days with viral loads too low to yield high quality sequences are shown by the gaps in data (i.e. cat 3 day 6 and cat 4 day 9).

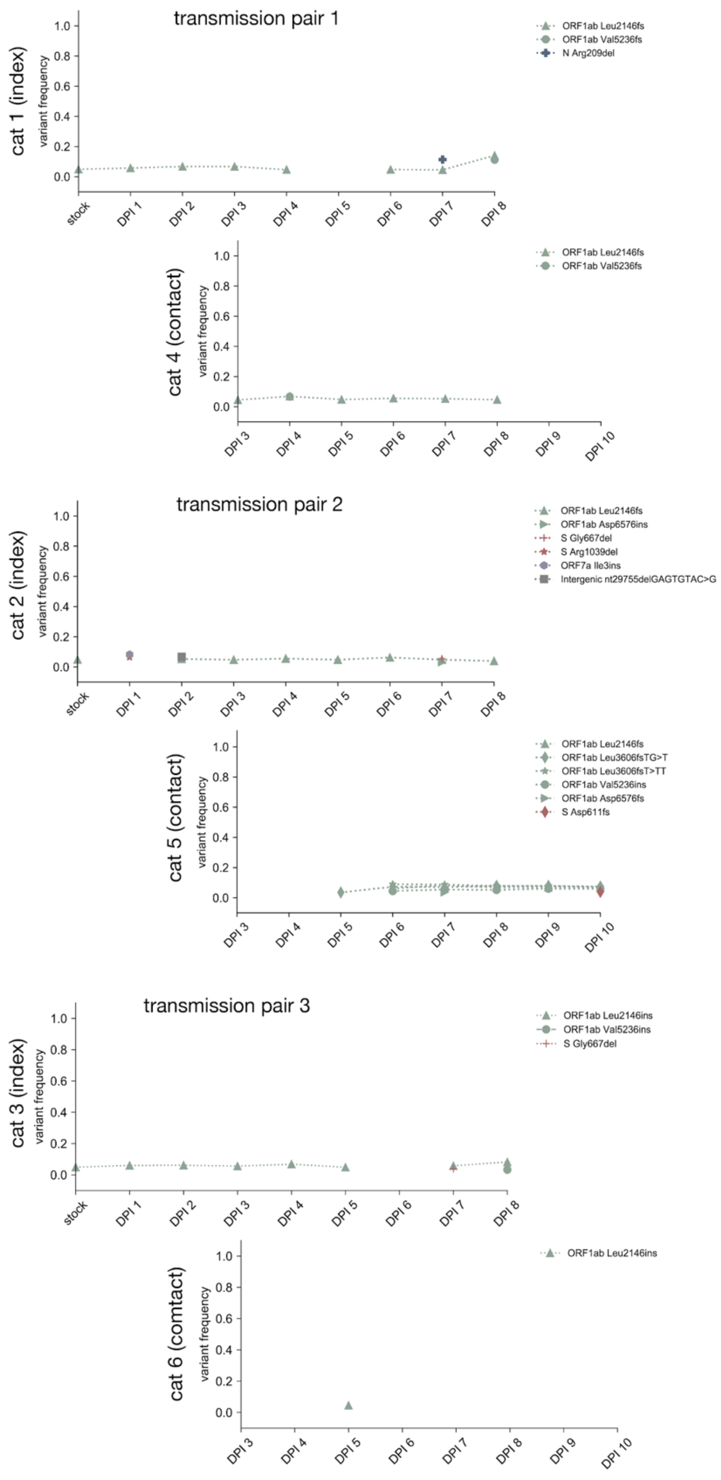


**Supplementary Figure 3. Longitudinal frequency of indels detected in all cats and at all timepoints.** Each indel is colored based on gene location. Days with viral loads too low to yield high quality sequences are shown by the gaps in data (i.e. cat 3 day 6 and cat 4 day 9).

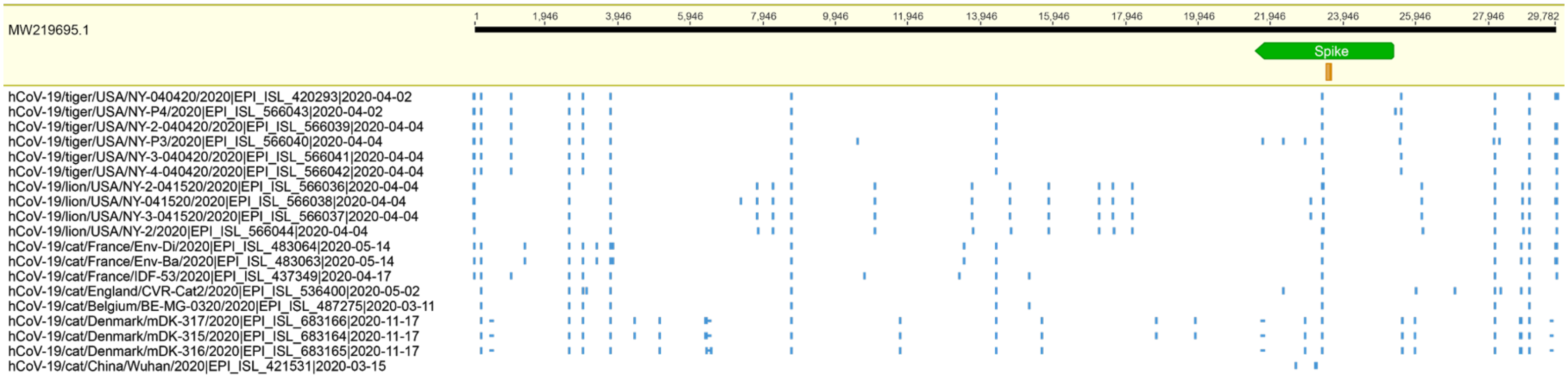


**Supplementary Figure 4. Sequence alignment of all tiger, lion, and domestic cat sequences available in GISAID as of December 2020.** Sequences were aligned against MW219695.1, the inoculum virus used in these experiments. Consensus-level differences are highlighted with a blue vertical marker. Indels are noted with a horizontal vertical marker. The spike open reading frame is annotated with a green marker and site amino acid 655 in Spike is highlighted with the orange box. None of these sequences contain a consensus mutation at residue 655 in Spike.

A

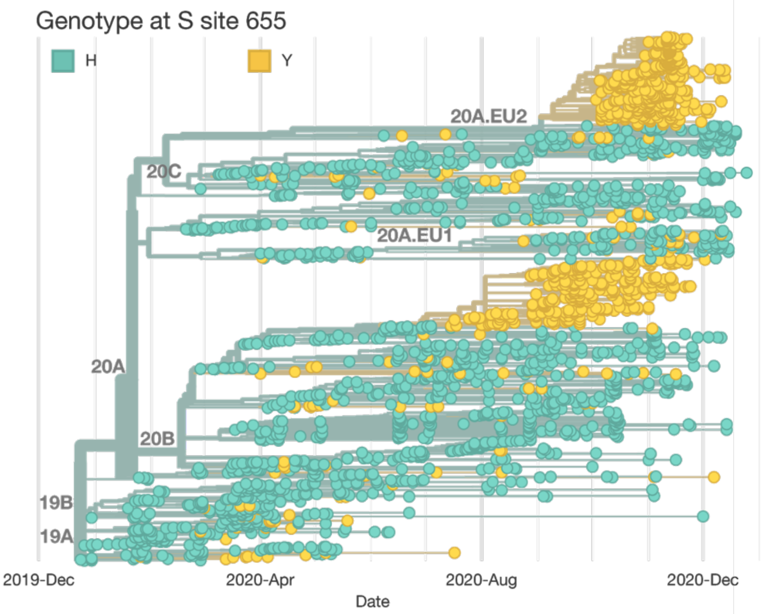


B

| Country | Date of first detected sequence with H655Y | Number of sequences | Date of most recent detected sequence with H655Y |
|---|---|---|---|
| United Kingdom | 2020-03-13 | 943 | 2020-12-13 |
| Denmark | 2020-04-13 | 44 | 2020-11-30 |
| USA | 2020-03-01 | 42 | 2020-11-12 |
| South Africa | 2020-06-11 | 8 | 2020-10-9 |
| India | 2020-05-11 | 6 | 2020-9-2 |
| China | 2020-01-26 | 3 | 2020-3-13 |
| Netherlands | 2020-05-07 | 3 | 2020-12-7 |
| Australia | 2020-07-31 | 3 | 2020-8-12 |
| Israel | 2020-03-24 | 2 | 2020-4-5 |
| United Arab Emirates | 2020-05-15 | 2 | 2020-5-18 |
| France | 2020-05-26 | 2 | 2020-8-14 |
| Rwanda | 2020-10-17 | 2 | 2020-10-22 |
| Taiwan | 2020-03-19 | 1 | 2020-3-19 |
| Portugal | 2020-04-02 | 1 | 2020-4-2 |
| Belize | 2020-04-09 | 1 | 2020-4-9 |
| Romania | 2020-04-09 | 1 | 2020-4-9 |
| Turkey | 2020-04-26 | 1 | 2020-4-26 |
| Brazil | 2020-06-30 | 1 | 2020-6-30 |
| Bangladesh | 2020-07-12 | 1 | 2020-7-12 |
| Czech Republic | 2020-11-04 | 1 | 2020-11-4 |
| Russia | 2020-11-16 | 1 | 2020-11-16 |
| Italy | 2020-12-02 | 1 | 2020-12-2 |

C

Geographic dispersion of SARS-CoV-2 viruses with Spike H655Y variant

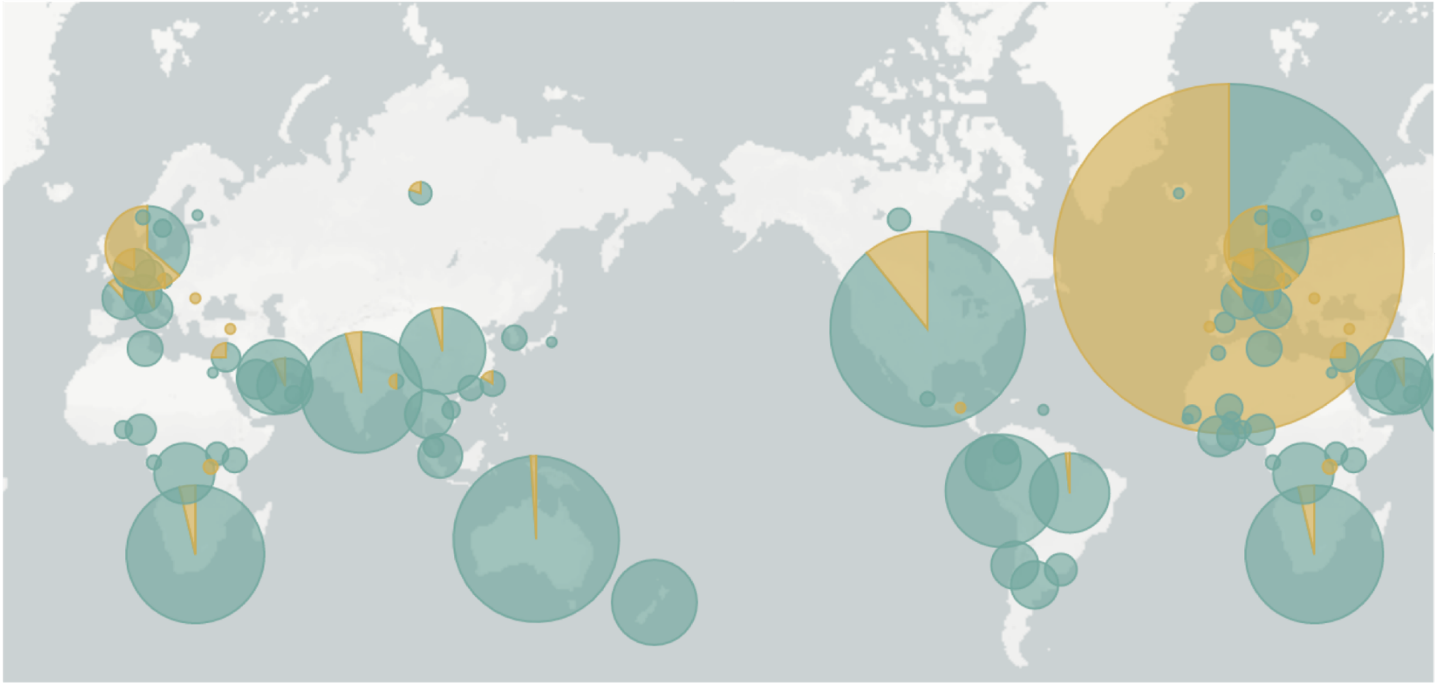

**Supplementary Figure 5. Geographic dispersion of Spike H655Y variant.** A) A time-resolved phylogeny focused on viruses that contain Spike H655Y. Viruses that contain histidine (H) at Spike 655 are colored in teal. Viruses with tyrosine (Y) at Spike 655 are colored in yellow. B) Counts of SARS-CoV-2 viruses that contain Spike H655Y, broken down by country. C) Map highlighting the number viruses from each country. The size of the circle represents the number of sequences from the appropriate country contained in the phylogeny.

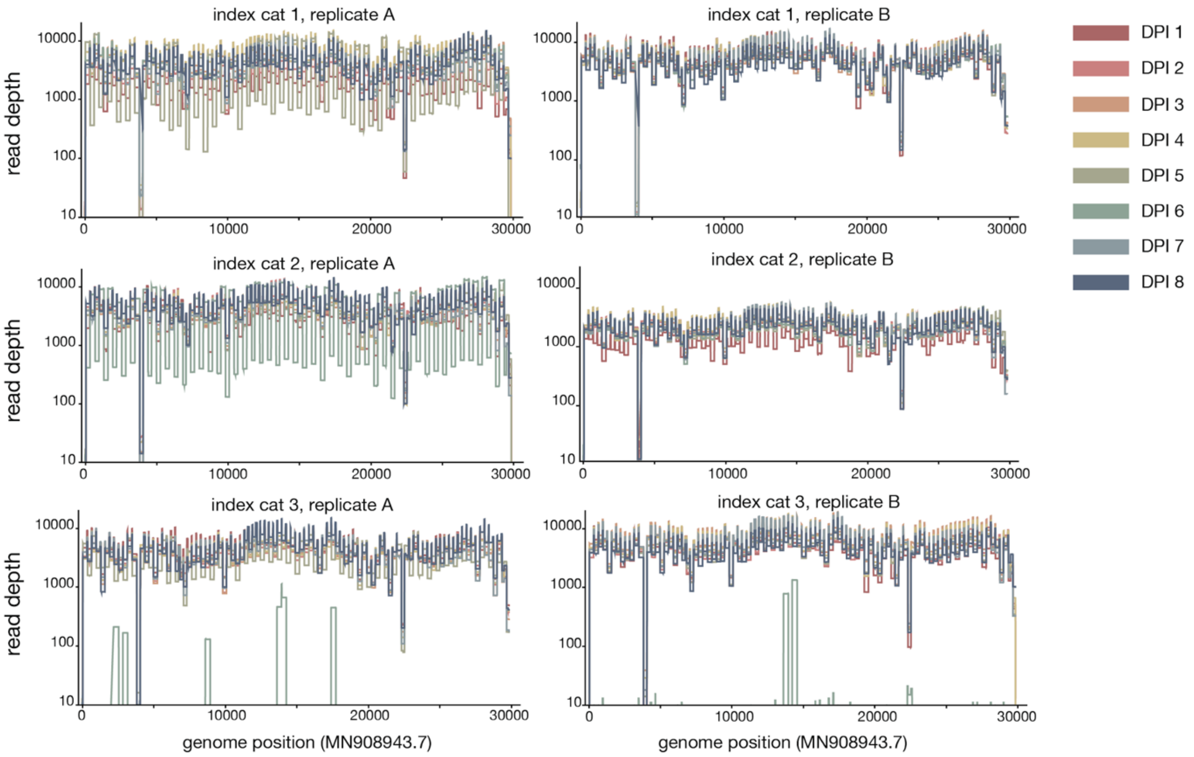


**Supplementary Figure 6. Read depth across the SARS-CoV-2 genome in index cats.** Each day is represented by a different color. Replicate A is shown in the left column and replicate B is shown in the right column.

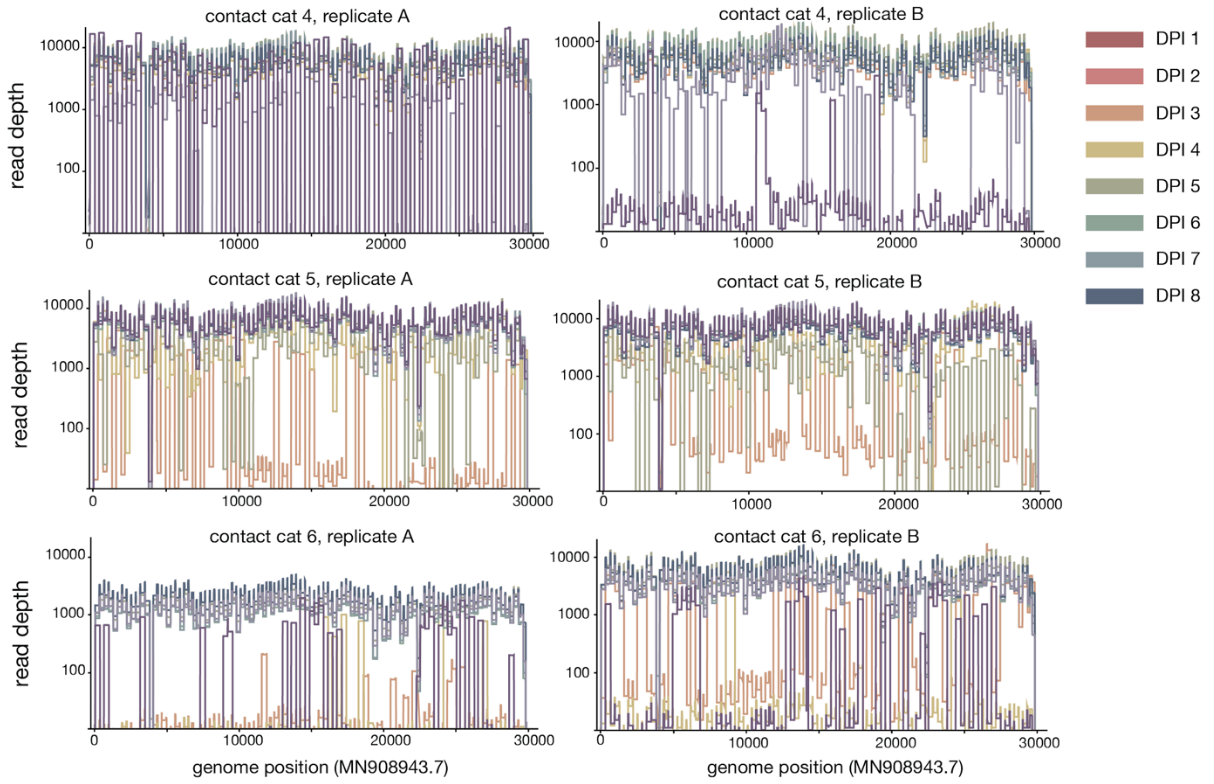


**Supplementary Figure 7. Read depth across the SARS-CoV-2 genome in contact cats.** Each day is represented by a different color. Replicate A is shown in the left column and replicate B is shown in the right column.

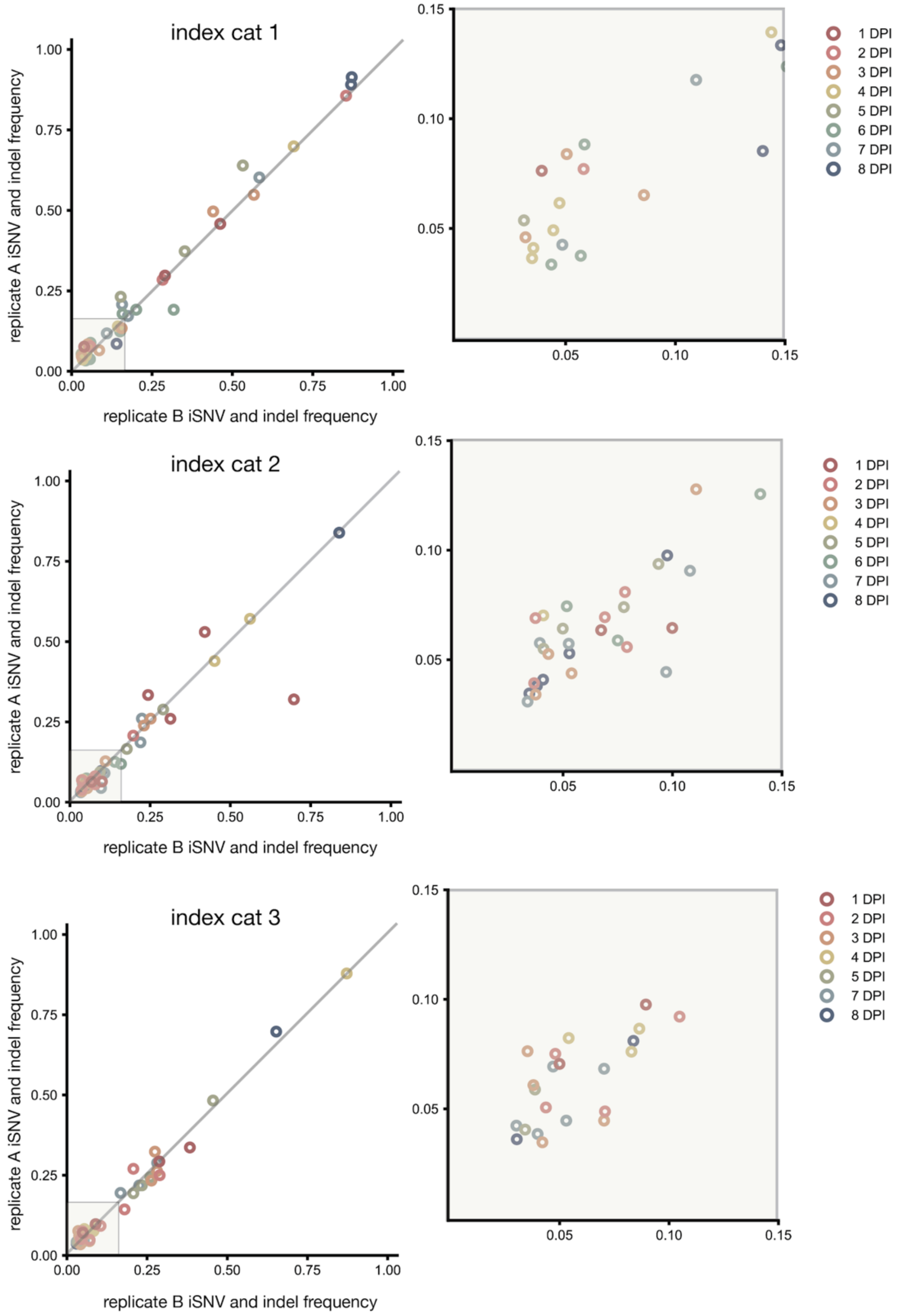


**Supplementary Figure 8. Intersection variants found across technical replicates in index cats.** The frequency of each variant per replicate is shown here. The diagonal line represents the 1:1 intersection of replicate variants. The subplot to the right of each primary plot is a zoomed-in view of the low-frequency variants (3-15%). Each timepoint is denoted by a different color.

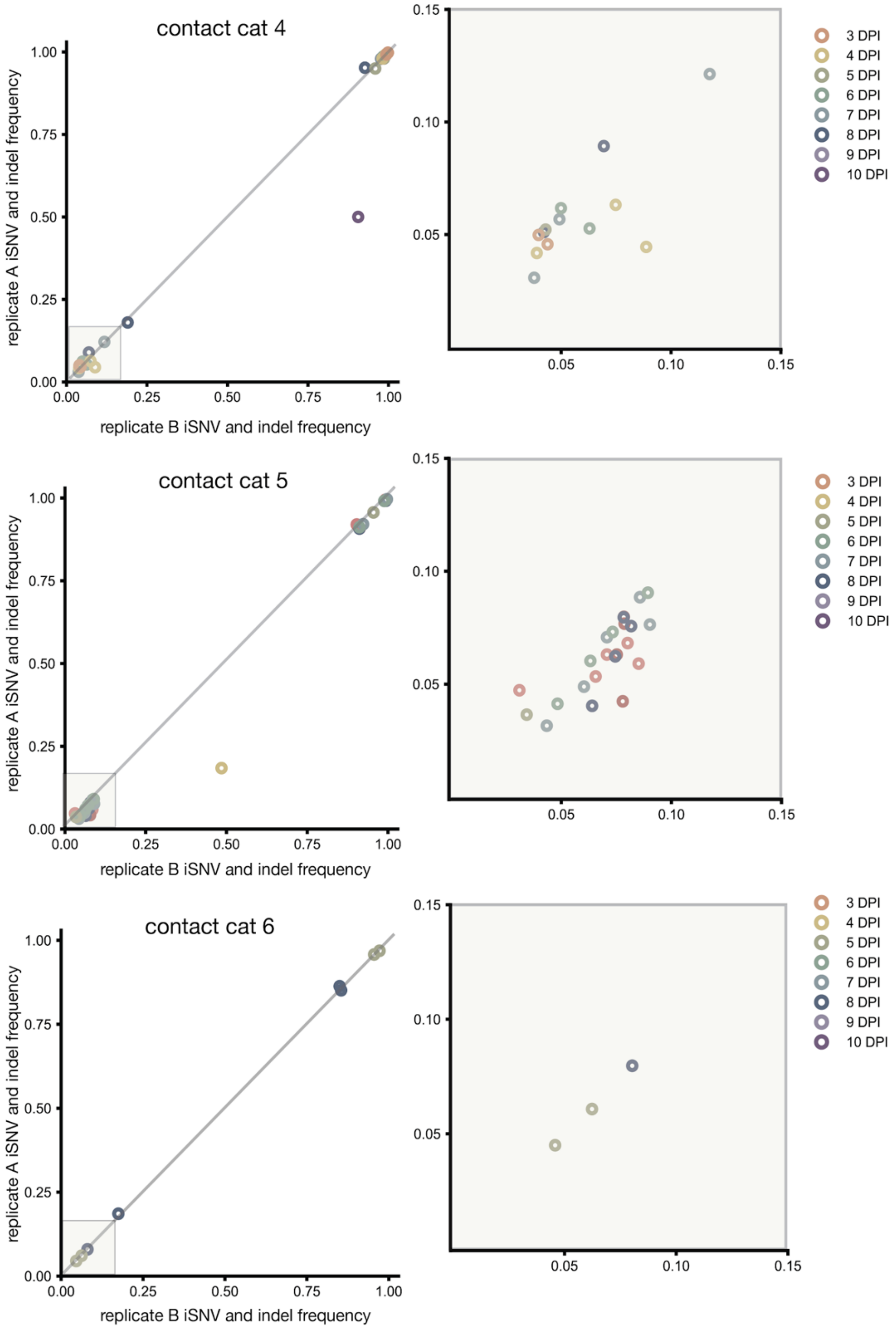


**Supplementary Figure 9. Intersection variants found across technical replicates in contact cats.** The frequency of each variant per replicate is shown here. The diagonal line represents the 1:1 intersection of replicate variants. The subplot to the right of each primary plot is a zoomed-in view of the low-frequency variants (3-15%). Each timepoint is denoted by a different color.

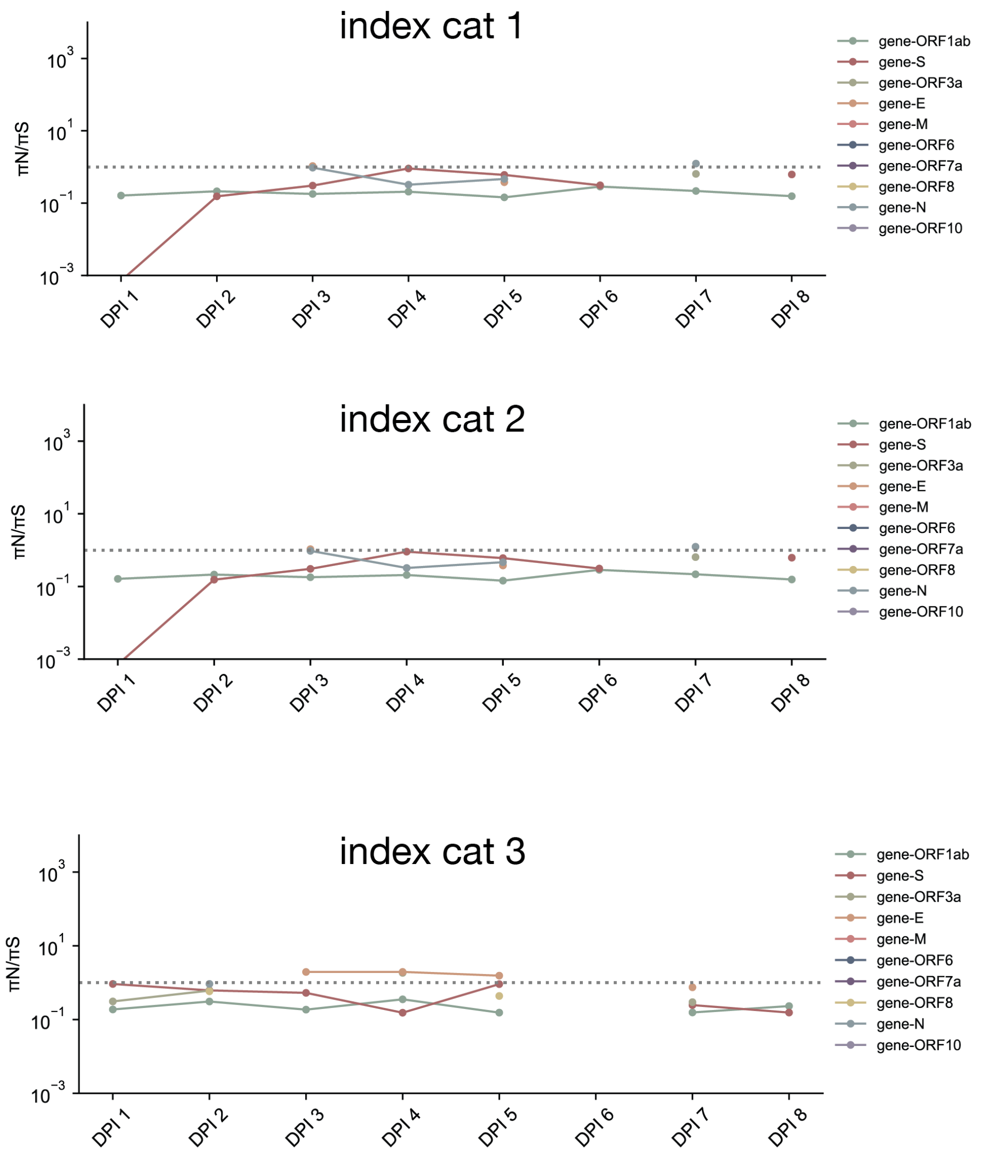


**Supplementary Figure 10. Longitudinal pairwise nonsynonymous nucleotide diversity divided by pairwise synonymous nucleotide diversity in index cats.** Line color denotes gene. The horizontal dotted gray line is plotted at $y = 1$ or when πN ~ πS.

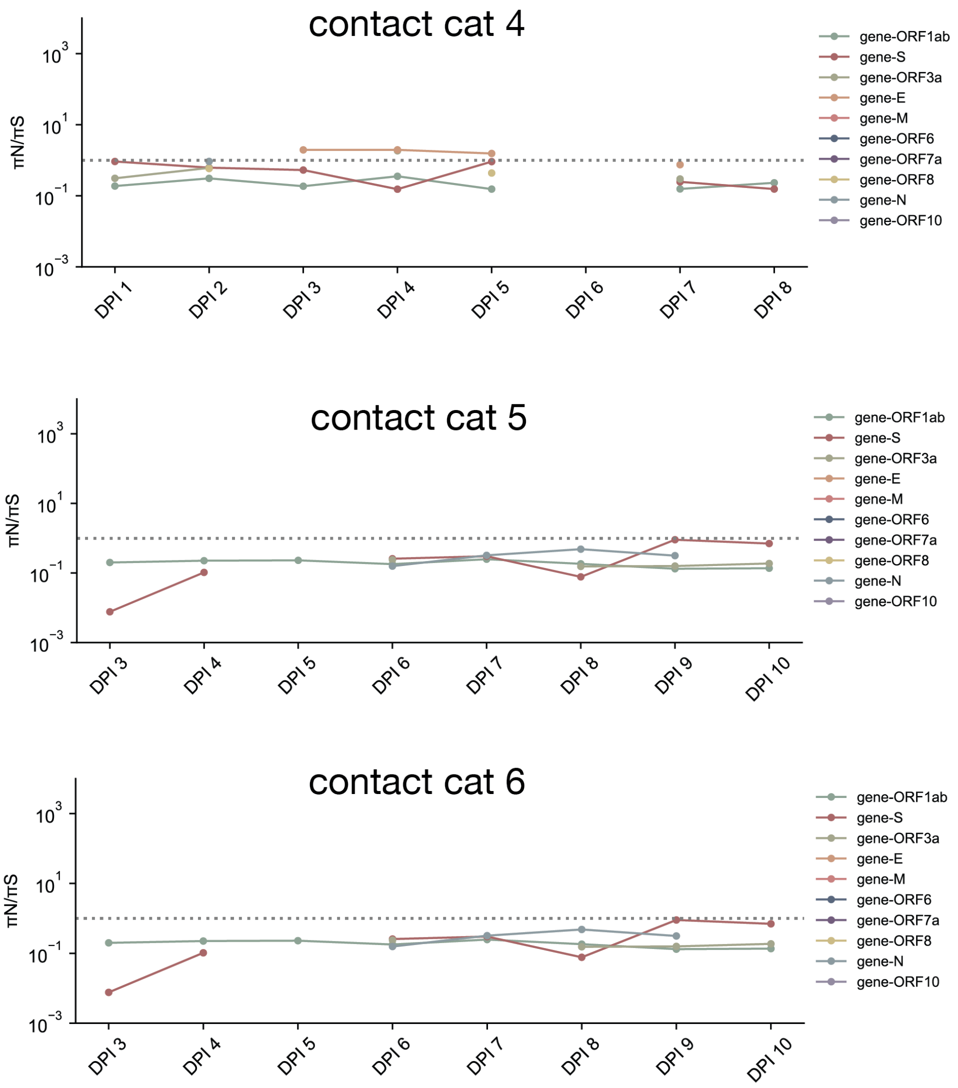


**Supplementary Figure 11. Longitudinal pairwise nonsynonymous nucleotide diversity divided by pairwise synonymous nucleotide diversity in contact cats.** Line color denotes gene. The horizontal dotted gray line is plotted at $y = 1$ or when $\pi N \sim \pi S$.

### Supplementary Tables

| | mean πS | std πS | mean πN | std πN | πN/πS | statistic, p-value |
|---|---|---|---|---|---|---|
| index cat 1 | | | | | | |
| **ORF1ab** | 0.015670 | 0.005609 | 0.003019 | 0.001124 | 0.195265 | (statistic=6.23, pvalue=2.11e-05) |
| **S** | 0.001564 | 0.000600 | 0.000644 | 0.000467 | 0.413995 | (statistic=3.34, pvalue=0.005) |
| **ORF3a** | 0.005367 | 0.000001 | 0.005878 | 0.002267 | 0.641899 | (statistic=-0.303, pvalue=0.772) |
| **E** | 0.011707 | 0.010139 | 0.011930 | 0.005719 | 0.719601 | (statistic=-0.0456, pvalue=0.965) |
| **M** | | | 0.002138 | 0.000059 | | (statistic=nan, pvalue=nan) |
| **ORF6** | | | 0.007395 | | | (statistic=nan, pvalue=nan) |
| **ORF7a** | 0.011992 | 0.000487 | | | | (statistic=nan, pvalue=nan) |
| **ORF8** | 0.031186 | 0.012756 | | | | (statistic=nan, pvalue=nan) |
| **N** | 0.005292 | 0.002941 | 0.003036 | 0.001403 | 0.744626 | (statistic=1.410, pvalue=0.196) |
| **ORF10** | | | | | | (statistic=nan, pvalue=nan) |
| index cat 2 | | | | | | |
| **ORF1ab** | 0.025570 | 0.007229 | 0.005423 | 0.001061 | 0.219071 | (statistic=7.799, pvalue=1.837e-06) |
| **S** | 0.004651 | 0.002567 | 0.001476 | 0.000556 | 0.457442 | (statistic=3.419, pvalue=0.0042) |
| **ORF3a** | 0.008660 | 0.003052 | 0.003867 | 0.001630 | 0.535879 | (statistic=3.549, pvalue=0.0053) |
| **E** | 0.008410 | 0.009696 | 0.015842 | 0.010396 | 436.372255 | (statistic=-1.191, pvalue=0.261) |
| **M** | | | 0.002479 | 0.000841 | | (statistic=nan, pvalue=nan) |
| **ORF6** | | | 0.007468 | | | (statistic=nan, pvalue=nan) |
| **ORF7a** | 0.011872 | | | | | (statistic=nan, pvalue=nan) |
| **ORF8** | 0.030343 | 0.012115 | 0.005673 | 0.002620 | 0.224566 | (statistic=2.733, pvalue=0.0292) |
| **N** | 0.006988 | 0.004962 | 0.001398 | 0.000535 | 0.346156 | (statistic=2.240, pvalue=0.066) |
| **ORF10** | | | | | | (statistic=nan, pvalue=nan) |
| index cat 3 | | | | | | |
| **ORF1ab** | 0.022195 | 0.005336 | 0.004752 | 0.000970 | 0.225357 | (statistic=8.509, pvalue=1.988e-06) |
| **S** | 0.003619 | 0.001697 | 0.001584 | 0.001032 | 0.505292 | (statistic=2.711, pvalue=0.0189) |
| **ORF3a** | 0.005516 | 0.000037 | 0.004351 | 0.003509 | 0.764893 | (statistic=0.649, pvalue=0.533) |
| **E** | 0.016174 | 0.000532 | 0.023985 | 0.007123 | 1.553581 | (statistic=-2.145, pvalue=0.0642) |
| **M** | 0.006492 | | 0.002078 | 0.000007 | | (statistic=nan, pvalue=nan) |
| **ORF6** | | | | | | (statistic=nan, pvalue=nan) |
| **ORF7a** | 0.012294 | 0.000305 | | | | (statistic=nan, pvalue=nan) |
| **ORF8** | 0.020154 | 0.006891 | 0.008740 | 0.002221 | 0.509847 | (statistic=2.708, pvalue=0.0352) |
| **N** | 0.005320 | 0.001974 | 0.003336 | | 0.925883 | (statistic=nan, pvalue=nan) |
| **ORF10** | | | | | | (statistic=nan, pvalue=nan) |

**Supplementary Table 1.** Nonsynonymous and synonymous nucleotide diversity estimates in index cats.

| | Cat 1 | Cat 2 | Cat 3 | Cat 4 | Cat 5 | Cat 6 |
|---|---|---|---|---|---|---|
| **π DPI 1** | 0.000246 | 0.000290 | 0.000433 | | | |
| **π DPI 2** | 0.000314 | 0.000705 | 0.000546 | | | |
| **π DPI 3** | 0.000458 | 0.000557 | 0.000781 | 0.000712 | 0.000153 | |
| **π DPI 4** | 0.000577 | 0.000650 | 0.000568 | 0.000796 | 0.000206 | 0.000037 |
| **π DPI 5** | 0.000489 | 0.000513 | 0.000540 | 0.001007 | 0.000149 | 0.000576 |
| **π DPI 6** | 0.000430 | 0.000720 | | 0.000917 | 0.000854 | 0.000156 |
| **π DPI 7** | 0.000365 | 0.000541 | 0.000683 | 0.000721 | 0.000876 | 0.000025 |
| **π DPI 8** | 0.000214 | 0.000591 | 0.000458 | 0.000879 | 0.000872 | 0.000720 |
| **π DPI 9** | | | | 0.000125 | 0.000965 | 0.000070 |
| **π DPI 10** | | | | 0.000371 | 0.000932 | 0.000000 |
| **mean π** | 0.000387 | 0.000571 | 0.000573 | 0.000691 | 0.000626 | 0.000226 |
| **std π** | 0.000117 | 0.000128 | 0.000114 | 0.000279 | 0.000355 | 0.000273 |

**Supplementary Table 2.** Genome-wide pairwise nucleotide diversity estimates in index and contact cats

| name | pool | sequence | length | %gc | tm (use 65) |
|---|---|---|---|---|---|
| nCoV-2019_1_LEFT | nCoV-2019_1 | ACCAACCAACTTTCGATCTCTTGT | 24 | 41.67 | 60.69 |
| nCoV-2019_1_RIGHT | nCoV-2019_1 | CATCTTTAAGATGTTGACGTGCCTC | 25 | 44 | 60.45 |
| nCoV-2019_2_LEFT | nCoV-2019_2 | CTGTTTTACAGGTTCGCGACGT | 22 | 50 | 61.67 |
| nCoV-2019_2_RIGHT | nCoV-2019_2 | TAAGGATCAGTGCCAAGCTCGT | 22 | 50 | 61.74 |
| nCoV-2019_3_LEFT | nCoV-2019_1 | CGGTAATAAAGGAGCTGGTGGC | 22 | 54.55 | 61.32 |
| nCoV-2019_3_RIGHT | nCoV-2019_1 | AAGGTGTCTGCAATTCATAGCTCT | 24 | 41.67 | 60.32 |
| nCoV-2019_4_LEFT | nCoV-2019_2 | GGTGTATACTGCTGCCGTGAAC | 22 | 54.55 | 61.56 |
| nCoV-2019_4_RIGHT | nCoV-2019_2 | CACAAGTAGTGGCACCTTCTTTAGT | 25 | 44 | 60.97 |
| nCoV-2019_5_LEFT | nCoV-2019_1 | TGGTGAAACTTCATGGCAGACG | 22 | 50 | 61.39 |
| nCoV-2019_5_RIGHT | nCoV-2019_1 | ATTGATGTTGACTTTCTCTTTTTGGAGT | 28 | 32.14 | 60.17 |
| nCoV-2019_6_LEFT | nCoV-2019_2 | GGTGTTGTTGGAGAAGGTTCCG | 22 | 54.55 | 61.64 |
| nCoV-2019_6_RIGHT | nCoV-2019_2 | TAGCGGCCTTCTGTAAAACACG | 22 | 50 | 61.18 |
| nCoV-2019_7_LEFT | nCoV-2019_1 | ATCAGAGGCTGCTCGTGTTGTA | 22 | 50 | 61.73 |
| nCoV-2019_7_LEFT_alt0 | nCoV-2019_1 | CATTTGCATCAGAGGCTGCTCG | 22 | 54.55 | 62.44 |
| nCoV-2019_7_RIGHT | nCoV-2019_1 | TGCACAGGTGACAATTTGTCCA | 22 | 45.45 | 60.95 |
| nCoV-2019_7_RIGHT_alt5 | nCoV-2019_1 | AGGTGACAATTTGTCCACCGAC | 22 | 50 | 61.07 |
| nCoV-2019_8_LEFT | nCoV-2019_2 | AGAGTTTCTTAGAGACGGTTGGGA | 24 | 45.83 | 61 |
| nCoV-2019_8_RIGHT | nCoV-2019_2 | GCTTCAACAGCTTCACTAGTAGGT | 24 | 45.83 | 60.56 |
| nCoV-2019_9_LEFT | nCoV-2019_1 | TCCCACAGAAGTGTTAACAGAGGA | 24 | 45.83 | 61.18 |
| nCoV-2019_9_LEFT_alt4 | nCoV-2019_1 | TTCCCACAGAAGTGTTAACAGAGG | 24 | 45.83 | 60.44 |
| nCoV-2019_9_RIGHT | nCoV-2019_1 | ATGACAGCATCTGCCACAACAC | 22 | 50 | 61.71 |
| nCoV-2019_9_RIGHT_alt2 | nCoV-2019_1 | GACAGCATCTGCCACAACACAG | 22 | 54.55 | 62.26 |
| nCoV-2019_10_LEFT | nCoV-2019_2 | TGAGAAGTGCTCTGCCTATACAGT | 24 | 45.83 | 61.12 |
| nCoV-2019_10_RIGHT | nCoV-2019_2 | TCATCTAACCAATCTTCTTCTTGCTCT | 27 | 37.04 | 60.31 |
| nCoV-2019_11_LEFT | nCoV-2019_1 | GGAATTTGGTGCCACTTCTGCT | 22 | 50 | 61.66 |
| nCoV-2019_11_RIGHT | nCoV-2019_1 | TCATCAGATTCAACTTGCATGGCA | 24 | 41.67 | 61.35 |
| nCoV-2019_12_LEFT | nCoV-2019_2 | AAACATGGAGGAGGTGTTGCAG | 22 | 50 | 61.08 |
| nCoV-2019_12_RIGHT | nCoV-2019_2 | TTCACTCTTCATTTCCAAAAAGCTTGA | 27 | 33.33 | 60.36 |
| nCoV-2019_13_LEFT | nCoV-2019_1 | TCGCACAAATGTCTACTTAGCTGT | 24 | 41.67 | 60.56 |
| nCoV-2019_13_RIGHT | nCoV-2019_1 | ACCACAGCAGTTAAAACACCCT | 22 | 45.45 | 60.36 |
| nCoV-2019_14_LEFT | nCoV-2019_2 | CATCCAGATTCTGCCACTCTTGT | 23 | 47.83 | 60.62 |

| | | | | | |
|---|---|---|---|---|---|
| nCoV-2019_14_LEFT_alt4 | nCoV-2019_2 | TGGCAATCTTCATCCAGATTCTGC | 24 | 45.83 | 61.47 |
| nCoV-2019_14_RIGHT | nCoV-2019_2 | AGTTTCCACACAGACAGGCATT | 22 | 45.45 | 60.42 |
| nCoV-2019_14_RIGHT_alt2 | nCoV-2019_2 | TGCGTGTTTCTTCTGCATGTGC | 22 | 50 | 62.76 |
| nCoV-2019_15_LEFT | nCoV-2019_1 | ACAGTGCTTAAAAAGTGTAAAAGTGCC | 27 | 37.04 | 61.32 |
| nCoV-2019_15_LEFT_alt1 | nCoV-2019_1 | AGTGCTTAAAAAGTGTAAAAGTGCCT | 26 | 34.62 | 60.13 |
| nCoV-2019_15_RIGHT | nCoV-2019_1 | AACAGAAACTGTAGCTGGCACT | 22 | 45.45 | 60.16 |
| nCoV-2019_15_RIGHT_alt3 | nCoV-2019_1 | ACTGTAGCTGGCACTTTGAGAGA | 23 | 47.83 | 61.57 |
| nCoV-2019_16_LEFT | nCoV-2019_2 | AATTTGGAAGAAGCTGCTCGGT | 22 | 45.45 | 60.82 |
| nCoV-2019_16_RIGHT | nCoV-2019_2 | CACAACTTGCGTGTGGAGGTTA | 22 | 50 | 61.32 |
| nCoV-2019_17_LEFT | nCoV-2019_1 | CTTCTTTCTTTGAGAGAAGTGAGGACT | 27 | 40.74 | 60.69 |
| nCoV-2019_17_RIGHT | nCoV-2019_1 | TTTGTTGGAGTGTTAACAATGCAGT | 25 | 36 | 60.11 |
| nCoV-2019_18_LEFT | nCoV-2019_2 | TGGAAATACCCACAAGTTAATGGTTTAAC | 29 | 34.48 | 60.69 |
| nCoV-2019_18_LEFT_alt2 | nCoV-2019_2 | ACTTCTATTAAATGGGCAGATAACAACTGT | 30 | 33.33 | 61.38 |
| nCoV-2019_18_RIGHT | nCoV-2019_2 | AGCTTGTTTACCACACGTACAAGG | 24 | 45.83 | 61.51 |
| nCoV-2019_18_RIGHT_alt1 | nCoV-2019_2 | GCTTGTTTACCACACGTACAAGG | 23 | 47.83 | 60.3 |
| nCoV-2019_19_LEFT | nCoV-2019_1 | GCTGTTATGTACATGGGCACACT | 23 | 47.83 | 61.18 |
| nCoV-2019_19_RIGHT | nCoV-2019_1 | TGTCCAACTTAGGGTCAATTTCTGT | 25 | 40 | 60.4 |
| nCoV-2019_20_LEFT | nCoV-2019_2 | ACAAAGAAAACAGTTACACAACAACCA | 27 | 33.33 | 60.68 |
| nCoV-2019_20_RIGHT | nCoV-2019_2 | ACGTGGCTTTATTAGTTGCATTGTT | 25 | 36 | 60.28 |
| nCoV-2019_21_LEFT | nCoV-2019_1 | TGGCTATTGATTATAAACACTACACACCC | 29 | 37.93 | 61.49 |
| nCoV-2019_21_LEFT_alt2 | nCoV-2019_1 | GGCTATTGATTATAAACACTACACACCCT | 29 | 37.93 | 61.29 |
| nCoV-2019_21_RIGHT | nCoV-2019_1 | TAGATCTGTGTGGCCAACCTCT | 22 | 50 | 60.83 |
| nCoV-2019_21_RIGHT_alt0 | nCoV-2019_1 | GATCTGTGTGGCCAACCTCTTC | 22 | 54.55 | 61.2 |
| nCoV-2019_22_LEFT | nCoV-2019_2 | ACTACCGAAGTTGTAGGAGACATTATACT | 29 | 37.93 | 61.25 |
| nCoV-2019_22_RIGHT | nCoV-2019_2 | ACAGTATTCTTTGCTATAGTAGTCGGC | 27 | 40.74 | 60.73 |
| nCoV-2019_23_LEFT | nCoV-2019_1 | ACAACTACTAACATAGTTACACGGTGT | 27 | 37.04 | 60.26 |
| nCoV-2019_23_RIGHT | nCoV-2019_1 | ACCAGTACAGTAGGTTGCAATAGTG | 25 | 44 | 60.57 |
| nCoV-2019_24_LEFT | nCoV-2019_2 | AGGCATGCCTTCTTACTGTACTG | 23 | 47.83 | 60.37 |
| nCoV-2019_24_RIGHT | nCoV-2019_2 | ACATTCTAACCATAGCTGAAATCGGG | 26 | 42.31 | 61.19 |
| nCoV-2019_25_LEFT | nCoV-2019_1 | GCAATTGTTTTTCAGCTATTTTGCAGT | 27 | 33.33 | 60.73 |
| nCoV-2019_25_RIGHT | nCoV-2019_1 | ACTGTAGTGACAAGTCTCTCGCA | 23 | 47.83 | 61.3 |
| nCoV-2019_26_LEFT | nCoV-2019_2 | TTGTGATACATTCTGTGCTGGTAGT | 25 | 40 | 60.28 |

| | | | | | |
|---|---|---|---|---|---|
| nCoV-2019_26_RIGHT | nCoV-2019_2 | TCCGCACTATCACCAACATCAG | 22 | 50 | 60.42 |
| nCoV-2019_27_LEFT | nCoV-2019_1 | ACTACAGTCAGCTTATGTGTCAACC | 25 | 44 | 60.8 |
| nCoV-2019_27_RIGHT | nCoV-2019_1 | AATACAAGCACCAAGGTCACGG | 22 | 50 | 61.13 |
| nCoV-2019_28_LEFT | nCoV-2019_2 | ACATAGAAGTTACTGGCGATAGTTGT | 26 | 38.46 | 60.13 |
| nCoV-2019_28_RIGHT | nCoV-2019_2 | TGTTTAGACATGACATGAACAGGTGT | 26 | 38.46 | 60.91 |
| nCoV-2019_29_LEFT | nCoV-2019_1 | ACTTGTGTTCCTTTTTGTTGCTGC | 24 | 41.67 | 61.39 |
| nCoV-2019_29_RIGHT | nCoV-2019_1 | AGTGTACTCTATAAGTTTTGATGGTGTGT | 29 | 34.48 | 60.69 |
| nCoV-2019_30_LEFT | nCoV-2019_2 | GCACAACTAATGGTGACTTTTTGCA | 25 | 40 | 61.19 |
| nCoV-2019_30_RIGHT | nCoV-2019_2 | ACCACTAGTAGATACACAAACACCAG | 26 | 42.31 | 60.3 |
| nCoV-2019_31_LEFT | nCoV-2019_1 | TTCTGAGTACTGTAGGCACGGC | 22 | 54.55 | 62.03 |
| nCoV-2019_31_RIGHT | nCoV-2019_1 | ACAGAATAAACACCAGGTAAGAATGAGT | 28 | 35.71 | 60.69 |
| nCoV-2019_32_LEFT | nCoV-2019_2 | TGGTGAATACAGTCATGTAGTTGCC | 25 | 44 | 61.09 |
| nCoV-2019_32_RIGHT | nCoV-2019_2 | AGCACATCACTACGCAACTTTAGA | 24 | 41.67 | 60.56 |
| nCoV-2019_33_LEFT | nCoV-2019_1 | ACTTTTGAAGAAGCTGCGCTGT | 22 | 45.45 | 61.58 |
| nCoV-2019_33_RIGHT | nCoV-2019_1 | TGGACAGTAAACTACGTCATCAAGC | 25 | 44 | 61.08 |
| nCoV-2019_34_LEFT | nCoV-2019_2 | TCCCATCTGGTAAAGTTGAGGGT | 23 | 47.83 | 61.02 |
| nCoV-2019_34_RIGHT | nCoV-2019_2 | AGTGAAATTGGGCCTCATAGCA | 22 | 45.45 | 60.03 |
| nCoV-2019_35_LEFT | nCoV-2019_1 | TGTTCGCATTCAACCAGGACAG | 22 | 50 | 61.39 |
| nCoV-2019_35_RIGHT | nCoV-2019_1 | ACTTCATAGCCACAAGGTTAAAGTCA | 26 | 38.46 | 60.69 |
| nCoV-2019_36_LEFT | nCoV-2019_2 | TTAGCTTGGTTGTACGCTGCTG | 22 | 50 | 61.44 |
| nCoV-2019_36_RIGHT | nCoV-2019_2 | GAACAAAGACCATTGAGTACTCTGGA | 26 | 42.31 | 60.74 |
| nCoV-2019_37_LEFT | nCoV-2019_1 | ACACACCACTGGTTGTTACTCAC | 23 | 47.83 | 60.93 |
| nCoV-2019_37_RIGHT | nCoV-2019_1 | GTCCACACTCTCCTAGCACCAT | 22 | 54.55 | 61.48 |
| nCoV-2019_38_LEFT | nCoV-2019_2 | ACTGTGTTATGTATGCATCAGCTGT | 25 | 40 | 60.86 |
| nCoV-2019_38_RIGHT | nCoV-2019_2 | CACCAAGAGTCAGTCTAAAGTAGCG | 25 | 48 | 61.13 |
| nCoV-2019_39_LEFT | nCoV-2019_1 | AGTATTGCCCTATTTTCTTCATAACTGGT | 29 | 34.48 | 61 |
| nCoV-2019_39_RIGHT | nCoV-2019_1 | TGTAACTGGACACATTGAGCCC | 22 | 50 | 60.55 |
| nCoV-2019_40_LEFT | nCoV-2019_2 | TGCACATCAGTAGTCTTACTCTCAGT | 26 | 42.31 | 61.25 |
| nCoV-2019_40_RIGHT | nCoV-2019_2 | CATGGCTGCATCACGGTCAAAT | 22 | 50 | 62.09 |
| nCoV-2019_41_LEFT | nCoV-2019_1 | GTTCCCTTCCATCATATGCAGCT | 23 | 47.83 | 60.75 |
| nCoV-2019_41_RIGHT | nCoV-2019_1 | TGGTATGACAACCATTAGTTTGGCT | 25 | 40 | 60.75 |
| nCoV-2019_42_LEFT | nCoV-2019_2 | TGCAAGAGATGGTTGTGTTCCC | 22 | 50 | 61.08 |

| | | | | | |
|---|---|---|---|---|---|
| nCoV-2019_42_RIGHT | nCoV-2019_2 | CCTACCTCCCTTTGTTGTGTTGT | 23 | 47.83 | 60.69 |
| nCoV-2019_43_LEFT | nCoV-2019_1 | TACGACAGATGTCTTGTGCTGC | 22 | 50 | 60.93 |
| nCoV-2019_43_RIGHT | nCoV-2019_1 | AGCAGCATCTACAGCAAAAGCA | 22 | 45.45 | 61.14 |
| nCoV-2019_44_LEFT | nCoV-2019_2 | TGCCACAGTACGTCTACAAGCT | 22 | 50 | 61.66 |
| nCoV-2019_44_LEFT_alt3 | nCoV-2019_2 | CCACAGTACGTCTACAAGCTGG | 22 | 54.55 | 60.67 |
| nCoV-2019_44_RIGHT | nCoV-2019_2 | AACCTTTCCACATACCGCAGAC | 22 | 50 | 60.87 |
| nCoV-2019_44_RIGHT_alt0 | nCoV-2019_2 | CGCAGACGGTACAGACTGTGTT | 22 | 54.55 | 62.77 |
| nCoV-2019_45_LEFT | nCoV-2019_1 | TACCTACAACTTGTGCTAATGACCC | 25 | 44 | 60.57 |
| nCoV-2019_45_LEFT_alt2 | nCoV-2019_1 | AGTATGTACAAATACCTACAACTTGTGCT | 29 | 34.48 | 60.94 |
| nCoV-2019_45_RIGHT | nCoV-2019_1 | AAATTGTTTCTTCATGTTGGTAGTTAGAGA | 30 | 30 | 60.01 |
| nCoV-2019_45_RIGHT_alt7 | nCoV-2019_1 | TTCATGTTGGTAGTTAGAGAAAGTGTGTC | 29 | 37.93 | 61.53 |
| nCoV-2019_46_LEFT | nCoV-2019_2 | TGTCGCTTCCAAGAAAAGGACG | 22 | 50 | 61.38 |
| nCoV-2019_46_LEFT_alt1 | nCoV-2019_2 | CGCTTCCAAGAAAAGGACGAAGA | 23 | 47.83 | 61.35 |
| nCoV-2019_46_RIGHT | nCoV-2019_2 | CACGTTCACCTAAGTTGGCGTA | 22 | 50 | 60.86 |
| nCoV-2019_46_RIGHT_alt2 | nCoV-2019_2 | CACGTTCACCTAAGTTGGCGTAT | 23 | 47.83 | 61.17 |
| nCoV-2019_47_LEFT | nCoV-2019_1 | AGGACTGGTATGATTTTGTAGAAAACCC | 28 | 39.29 | 61.42 |
| nCoV-2019_47_RIGHT | nCoV-2019_1 | AATAACGGTCAAAGAGTTTTAACCTCTC | 28 | 35.71 | 60.06 |
| nCoV-2019_48_LEFT | nCoV-2019_2 | TGTTGACACTGACTTAACAAAGCCT | 25 | 40 | 61.09 |
| nCoV-2019_48_RIGHT | nCoV-2019_2 | TAGATTACCAGAAGCAGCGTGC | 22 | 50 | 60.74 |
| nCoV-2019_49_LEFT | nCoV-2019_1 | AGGAATTACTTGTGTATGCTGCTGA | 25 | 40 | 60.57 |
| nCoV-2019_49_RIGHT | nCoV-2019_1 | TGACGATGACTTGGTTAGCATTAATACA | 28 | 35.71 | 61.05 |
| nCoV-2019_50_LEFT | nCoV-2019_2 | GTTGATAAGTACTTTGATTGTTACGATGGT | 30 | 33.33 | 60.59 |
| nCoV-2019_50_RIGHT | nCoV-2019_2 | TAACATGTTGTGCCAACCACCA | 22 | 45.45 | 60.95 |
| nCoV-2019_51_LEFT | nCoV-2019_1 | TCAATAGCCGCCACTAGAGGAG | 22 | 54.55 | 61.34 |
| nCoV-2019_51_RIGHT | nCoV-2019_1 | AGTGCATTAACATTGGCCGTGA | 22 | 45.45 | 61.14 |
| nCoV-2019_52_LEFT | nCoV-2019_2 | CATCAGGAGATGCCACAACTGC | 22 | 54.55 | 61.83 |
| nCoV-2019_52_RIGHT | nCoV-2019_2 | GTTGAGAGCAAAATTCATGAGGTCC | 25 | 44 | 60.62 |
| nCoV-2019_53_LEFT | nCoV-2019_1 | AGCAAAATGTTGGACTGAGACTGA | 24 | 41.67 | 60.69 |
| nCoV-2019_53_RIGHT | nCoV-2019_1 | AGCCTCATAAAACTCAGGTTCCC | 23 | 47.83 | 60.31 |
| nCoV-2019_54_LEFT | nCoV-2019_2 | TGAGTTAACAGGACACATGTTAGACA | 26 | 38.46 | 60.18 |
| nCoV-2019_54_RIGHT | nCoV-2019_2 | AACCAAAAACTTGTCCATTAGCACA | 25 | 36 | 60.11 |
| nCoV-2019_55_LEFT | nCoV-2019_1 | ACTCAACTTTACTTAGGAGGTATGAGCT | 28 | 39.29 | 61.43 |

| | | | | | |
|---|---|---|---|---|---|
| nCoV-2019_55_RIGHT | nCoV-2019_1 | GGTGTACTCTCCTATTTGTACTTTACTGT | 29 | 37.93 | 60.54 |
| nCoV-2019_56_LEFT | nCoV-2019_2 | ACCTAGACCACCACTTAACCGA | 22 | 50 | 60.49 |
| nCoV-2019_56_RIGHT | nCoV-2019_2 | ACACTATGCGAGCAGAAGGGTA | 22 | 50 | 61.21 |
| nCoV-2019_57_LEFT | nCoV-2019_1 | ATTCTACACTCCAGGGACCACC | 22 | 54.55 | 61.16 |
| nCoV-2019_57_RIGHT | nCoV-2019_1 | GTAATTGAGCAGGGTCGCCAAT | 22 | 50 | 61.26 |
| nCoV-2019_58_LEFT | nCoV-2019_2 | TGATTTGAGTGTTGTCAATGCCAGA | 25 | 40 | 61.44 |
| nCoV-2019_58_RIGHT | nCoV-2019_2 | CTTTTCTCCAAGCAGGGTTACGT | 23 | 47.83 | 61.06 |
| nCoV-2019_59_LEFT | nCoV-2019_1 | TCACGCATGATGTTTCATCTGCA | 23 | 43.48 | 61.42 |
| nCoV-2019_59_RIGHT | nCoV-2019_1 | AAGAGTCCTGTTACATTTTCAGCTTG | 26 | 38.46 | 60.02 |
| nCoV-2019_60_LEFT | nCoV-2019_2 | TGATAGAGACCTTTATGACAAGTTGCA | 27 | 37.04 | 60.53 |
| nCoV-2019_60_RIGHT | nCoV-2019_2 | GGTACCAACAGCTTCTCTAGTAGC | 24 | 50 | 60.44 |
| nCoV-2019_61_LEFT | nCoV-2019_1 | TGTTTATCACCCGCGAAGAAGC | 22 | 50 | 61.5 |
| nCoV-2019_61_RIGHT | nCoV-2019_1 | ATCACATAGACAACAGGTGCGC | 22 | 50 | 61.25 |
| nCoV-2019_62_LEFT | nCoV-2019_2 | GGCACATGGCTTTGAGTTGACA | 22 | 50 | 61.91 |
| nCoV-2019_62_RIGHT | nCoV-2019_2 | GTTGAACCTTTCTACAAGCCGC | 22 | 50 | 60.35 |
| nCoV-2019_63_LEFT | nCoV-2019_1 | TGTTAAGCGTGTTGACTGGACT | 22 | 45.45 | 60.16 |
| nCoV-2019_63_RIGHT | nCoV-2019_1 | ACAAACTGCCACCATCACAACC | 22 | 50 | 61.85 |
| nCoV-2019_64_LEFT | nCoV-2019_2 | TCGATAGATATCCTGCTAATTCCATTGT | 28 | 35.71 | 60.11 |
| nCoV-2019_64_RIGHT | nCoV-2019_2 | AGTCTTGTAAAAGTGTTCCAGAGGT | 25 | 40 | 60.1 |
| nCoV-2019_65_LEFT | nCoV-2019_1 | GCTGGCTTTAGCTTGTGGGTTT | 22 | 50 | 61.92 |
| nCoV-2019_65_RIGHT | nCoV-2019_1 | TGTCAGTCATAGAACAAACACCAATAGT | 28 | 35.71 | 60.9 |
| nCoV-2019_66_LEFT | nCoV-2019_2 | GGGTGTGGACATTGCTGCTAAT | 22 | 50 | 61.21 |
| nCoV-2019_66_RIGHT | nCoV-2019_2 | TCAATTTCCATTTGACTCCTGGGT | 24 | 41.67 | 60.45 |
| nCoV-2019_67_LEFT | nCoV-2019_1 | GTTGTCCAACAATTACCTGAAACTTACT | 28 | 35.71 | 60.43 |
| nCoV-2019_67_RIGHT | nCoV-2019_1 | CAACCTTAGAAACTACAGATAAATCTTGGG | 30 | 36.67 | 60.4 |
| nCoV-2019_68_LEFT | nCoV-2019_2 | ACAGGTTCATCTAAGTGTGTGTGT | 24 | 41.67 | 60.14 |
| nCoV-2019_68_RIGHT | nCoV-2019_2 | CTCCTTTATCAGAACCAGCACCA | 23 | 47.83 | 60.31 |
| nCoV-2019_69_LEFT | nCoV-2019_1 | TGTCGCAAAATATACTCAACTGTGTCA | 27 | 37.04 | 61.43 |
| nCoV-2019_69_RIGHT | nCoV-2019_1 | TCTTTATAGCCACGGAACCTCCA | 23 | 47.83 | 61.14 |
| nCoV-2019_70_LEFT | nCoV-2019_2 | ACAAAAGAAAATGACTCTAAAGAGGGTTT | 29 | 31.03 | 60.13 |
| nCoV-2019_70_RIGHT | nCoV-2019_2 | TGACCTTCTTTTAAAGACATAACAGCAG | 28 | 35.71 | 60.27 |
| nCoV-2019_71_LEFT | nCoV-2019_1 | ACAAATCCAATTCAGTTGTCTTCCTATTC | 29 | 34.48 | 60.54 |

| | | | | | |
|---|---|---|---|---|---|
| nCoV-2019_71_RIGHT | nCoV-2019_1 | TGGAAAAGAAAGGTAAGAACAAGTCCT | 27 | 37.04 | 60.8 |
| nCoV-2019_72_LEFT | nCoV-2019_2 | ACACGTGGTGTTTATTACCCTGAC | 24 | 45.83 | 61.04 |
| nCoV-2019_72_RIGHT | nCoV-2019_2 | ACTCTGAACTCACTTTCCATCCAAC | 25 | 44 | 60.97 |
| nCoV-2019_73_LEFT | nCoV-2019_1 | CAATTTTGTAATGATCCATTTTTGGGTGT | 29 | 31.03 | 60.29 |
| nCoV-2019_73_RIGHT | nCoV-2019_1 | CACCAGCTGTCCAACCTGAAGA | 22 | 54.55 | 62.45 |
| nCoV-2019_74_LEFT | nCoV-2019_2 | ACATCACTAGGTTTCAAACTTTACTTGC | 28 | 35.71 | 60.68 |
| nCoV-2019_74_RIGHT | nCoV-2019_2 | GCAACACAGTTGCTGATTCTCTTC | 24 | 45.83 | 60.85 |
| nCoV-2019_75_LEFT | nCoV-2019_1 | AGAGTCCAACCAACAGAATCTATTGT | 26 | 38.46 | 60.24 |
| nCoV-2019_75_RIGHT | nCoV-2019_1 | ACCACCAACCTTAGAATCAAGATTGT | 26 | 38.46 | 60.69 |
| nCoV-2019_76_LEFT | nCoV-2019_2 | AGGGCAAACTGGAAAGATTGCT | 22 | 45.45 | 60.76 |
| nCoV-2019_76_LEFT_alt3 | nCoV-2019_2 | GGGCAAACTGGAAAGATTGCTGA | 23 | 47.83 | 61.87 |
| nCoV-2019_76_RIGHT | nCoV-2019_2 | ACACCTGTGCCTGTTAAACCAT | 22 | 45.45 | 60.42 |
| nCoV-2019_76_RIGHT_alt0 | nCoV-2019_2 | ACCTGTGCCTGTTAAACCATTGA | 23 | 43.48 | 60.69 |
| nCoV-2019_77_LEFT | nCoV-2019_1 | CCAGCAACTGTTTGTGGACCTA | 22 | 50 | 60.75 |
| nCoV-2019_77_RIGHT | nCoV-2019_1 | CAGCCCCTATTAAACAGCCTGC | 22 | 54.55 | 61.59 |
| nCoV-2019_78_LEFT | nCoV-2019_2 | CAACTTACTCCTACTTGGCGTGT | 23 | 47.83 | 60.55 |
| nCoV-2019_78_RIGHT | nCoV-2019_2 | TGTGTACAAAAACTGCCATATTGCA | 25 | 36 | 60.22 |
| nCoV-2019_79_LEFT | nCoV-2019_1 | GTGGTGATTCAACTGAATGCAGC | 23 | 47.83 | 60.92 |
| nCoV-2019_79_RIGHT | nCoV-2019_1 | CATTTCATCTGTGAGCAAAGGTGG | 24 | 45.83 | 60.62 |
| nCoV-2019_80_LEFT | nCoV-2019_2 | TTGCCTTGGTGATATTGCTGCT | 22 | 45.45 | 60.89 |
| nCoV-2019_80_RIGHT | nCoV-2019_2 | TGGAGCTAAGTTGTTTAACAAGCG | 24 | 41.67 | 60.02 |
| nCoV-2019_81_LEFT | nCoV-2019_1 | GCACTTGGAAAACTTCAAGATGTGG | 25 | 44 | 61.24 |
| nCoV-2019_81_RIGHT | nCoV-2019_1 | GTGAAGTTCTTTTCTTGTGCAGGG | 24 | 45.83 | 60.73 |
| nCoV-2019_82_LEFT | nCoV-2019_2 | GGGCTATCATCTTATGTCCTTCCCT | 25 | 48 | 61.52 |
| nCoV-2019_82_RIGHT | nCoV-2019_2 | TGCCAGAGATGTCACCTAAATCAA | 24 | 41.67 | 60.02 |
| nCoV-2019_83_LEFT | nCoV-2019_1 | TCCTTTGCAACCTGAATTAGACTCA | 25 | 40 | 60.46 |
| nCoV-2019_83_RIGHT | nCoV-2019_1 | TTTGACTCCTTTGAGCACTGGC | 22 | 50 | 61.33 |
| nCoV-2019_84_LEFT | nCoV-2019_2 | TGCTGTAGTTGTCTCAAGGGCT | 22 | 50 | 61.61 |
| nCoV-2019_84_RIGHT | nCoV-2019_2 | AGGTGTGAGTAAACTGTTACAAACAAC | 27 | 37.04 | 60.36 |
| nCoV-2019_85_LEFT | nCoV-2019_1 | ACTAGCACTCTCCAAGGGTGTT | 22 | 50 | 61.03 |
| nCoV-2019_85_RIGHT | nCoV-2019_1 | ACACAGTCTTTTACTCCAGATTCCC | 25 | 44 | 60.51 |
| nCoV-2019_86_LEFT | nCoV-2019_2 | TCAGGTGATGGCACAACAAGTC | 22 | 50 | 61.07 |

| | | | | | |
|---|---|---|---|---|---|
| nCoV-2019_86_RIGHT | nCoV-2019_2 | ACGAAAGCAAGAAAAAGAAGTACGC | 25 | 40 | 61.01 |
| nCoV-2019_87_LEFT | nCoV-2019_1 | CGACTACTAGCGTGCCTTTGTA | 22 | 50 | 60.16 |
| nCoV-2019_87_RIGHT | nCoV-2019_1 | ACTAGGTTCCATTGTTCAAGGAGC | 24 | 45.83 | 60.81 |
| nCoV-2019_88_LEFT | nCoV-2019_2 | CCATGGCAGATTCCAACGGTAC | 22 | 54.55 | 61.58 |
| nCoV-2019_88_RIGHT | nCoV-2019_2 | TGGTCAGAATAGTGCCATGGAGT | 23 | 47.83 | 61.4 |
| nCoV-2019_89_LEFT | nCoV-2019_1 | GTACGCGTTCCATGTGGTCATT | 22 | 50 | 61.5 |
| nCoV-2019_89_LEFT_alt2 | nCoV-2019_1 | CGCGTTCCATGTGGTCATTCAA | 22 | 50 | 62.01 |
| nCoV-2019_89_RIGHT | nCoV-2019_1 | ACCTGAAAGTCAACGAGATGAAACA | 25 | 40 | 60.91 |
| nCoV-2019_89_RIGHT_alt4 | nCoV-2019_1 | ACGAGATGAAACATCTGTTGTCACT | 25 | 40 | 60.74 |
| nCoV-2019_90_LEFT | nCoV-2019_2 | ACACAGACCATTCCAGTAGCAGT | 23 | 47.83 | 61.58 |
| nCoV-2019_90_RIGHT | nCoV-2019_2 | TGAAATGGTGAATTGCCCTCGT | 22 | 45.45 | 60.82 |
| nCoV-2019_91_LEFT | nCoV-2019_1 | TCACTACCAAGAGTGTGTTAGAGGT | 25 | 44 | 60.93 |
| nCoV-2019_91_RIGHT | nCoV-2019_1 | TTCAAGTGAGAACCAAAAGATAATAAGCA | 29 | 31.03 | 60.03 |
| nCoV-2019_92_LEFT | nCoV-2019_2 | TTTGTGCTTTTTAGCCTTTCTGCT | 24 | 37.5 | 60.14 |
| nCoV-2019_92_RIGHT | nCoV-2019_2 | AGGTTCCTGGCAATTAATTGTAAAAGG | 27 | 37.04 | 60.53 |
| nCoV-2019_93_LEFT | nCoV-2019_1 | TGAGGCTGGTTCTAAATCACCCA | 23 | 47.83 | 61.59 |
| nCoV-2019_93_RIGHT | nCoV-2019_1 | AGGTCTTCCTTGCCATGTTGAG | 22 | 50 | 60.55 |
| nCoV-2019_94_LEFT | nCoV-2019_2 | GGCCCCAAGGTTTACCCAATAA | 22 | 50 | 60.56 |
| nCoV-2019_94_RIGHT | nCoV-2019_2 | TTTGGCAATGTTGTTCCTTGAGG | 23 | 43.48 | 60.18 |
| nCoV-2019_95_LEFT | nCoV-2019_1 | TGAGGGAGCCTTGAATACACCA | 22 | 50 | 61.1 |
| nCoV-2019_95_RIGHT | nCoV-2019_1 | CAGTACGTTTTTGCCGAGGCTT | 22 | 50 | 61.95 |
| nCoV-2019_96_LEFT | nCoV-2019_2 | GCCAACAACAACAAGGCCAAAC | 22 | 50 | 61.82 |
| nCoV-2019_96_RIGHT | nCoV-2019_2 | TAGGCTCTGTTGGTGGGAATGT | 22 | 50 | 61.36 |
| nCoV-2019_97_LEFT | nCoV-2019_1 | TGGATGACAAAGATCCAAATTTCAAAGA | 28 | 32.14 | 60.22 |
| nCoV-2019_97_RIGHT | nCoV-2019_1 | ACACACTGATTAAAGATTGCTATGTGAG | 28 | 35.71 | 60.17 |
| nCoV-2019_98_LEFT | nCoV-2019_2 | AACAATTGCAACAATCCATGAGCA | 24 | 37.5 | 60.5 |
| nCoV-2019_98_RIGHT | nCoV-2019_2 | TTCTCCTAAGAAGCTATTAAAATCACATGG | 30 | 33.33 | 60.01 |

**Supplementary Table 3. ARTIC v3 primer sequences.**
